# Integrative inference of subclonal tumour evolution from single-cell and bulk sequencing data

**DOI:** 10.1101/234914

**Authors:** Salem Malikic, Katharina Jahn, Jack Kuipers, S. Cenk Sahinalp, Niko Beerenwinkel

**Author notes:** Join first authors (Salem Malikic, Katharina Jahn, Jack Kuipers). Joint last and corresponding authors.

## Abstract

Understanding the evolutionary history and subclonal composition of a tumour represents one of the key challenges in overcoming treatment failure due to resistant cell populations. Most of the current data on tumour genetics stems from short read bulk sequencing data. While this type of data is characterised by low sequencing noise and cost, it consists of aggregate measurements across a large number of cells. It is therefore of limited use for the accurate detection of the distinct cellular populations present in a tumour and the unambiguous inference of their evolutionary relationships. Single-cell DNA sequencing instead provides data of the highest resolution for studying intra-tumour heterogeneity and evolution, but is characterised by higher sequencing costs and elevated noise rates. In this work, we develop the first computational approach that infers trees of tumour evolution from combined single-cell and bulk sequencing data. Using a comprehensive set of simulated data, we show that our approach systematically outperforms existing methods with respect to tree reconstruction accuracy and subclone identification. High fidelity reconstructions are obtained even with a modest number of single cells. We also show that combining single-cell and bulk sequencing data provides more realistic mutation histories for real tumours.

## 1 Introduction

Cancer is a genetic disease that develops through a branched evolutionary process [1]. It is characterised by the emergence of genetically distinct subclones through the random acquisition of mutations at the level of single-cells and shifting prevalences at the subclone level through selective advantages purveyed by driver mutations. This interplay creates complex mixtures of tumour cell populations which exhibit different susceptibility to targeted cancer therapies and are suspected to be the cause of treatment failure [2, 3]. Therefore it is of great interest to obtain a better understanding of the evolutionary histories of individual tumours and their subclonal composition [4].

Most genetic analyses of tumours are presently based on next-generation sequencing data of bulk tumour samples. Such data provide indirect measurements of the subclonal tumour composition in form of aggregate total and variant read counts measured across hundred thousands or millions of cells. A large number of approaches have been published over the last years that try to identify subclones, their frequencies, and in some cases, their phylogenetic relationships by deconvolving these aggregate data [5, 6, 7, 8, 9, 10, 11, 12, 13, 14, 15, 16, 17, 18, 19, 20, 21, 22, 23]. However, the underlying statistical problem is underdetermined [24, 25], and mutations with similar VAFs are automatically clustered into a single subclone. This inevitably leads to incorrect phylogenies for tumours with multiple distinct subclones of similar prevalences. The aggregate sequencing data additionally poses a limitation to the achievable tree resolution, as mutational signals of smaller subclones can not be distinguished from noise [26] and therefore not be reliably represented in the tree. Sequencing multiple samples from the same tumour and increasing the coverage can to some extent mitigate these issues but is not always practicable.

Another solution is the use of single-cell sequencing (SCS) data which directly provides mutation profiles of individual cells such that the phylogeny can be directly inferred without any form of deconvolution. The main challenge here instead are the high levels of noise found in SCS data that are primarily introduced during DNA amplification, a necessary step to obtain sufficient DNA material for sequencing. False negatives are the most prevalent error type due to allelic dropout, but also false positives occur when an error is introduced early in the amplification. Further noise stems from doublet mutation profiles, which occur when two cells are accidentally sequenced together [27]. Classic approaches for phylogeny reconstruction are not suitable for dealing with these SCS specific noise profiles, and a number of probabilistic approaches have been developed to specifically account for the error types found in SCS data [28, 29, 30, 31, 32].

A major difference between the evolutionary histories of tumours inferred from bulk and SCS data is that the former typically are clonal trees where mutations with similar frequencies are clustered together and represented in a single tree node (Figure 1a), while trees derived from SCS data are fully resolved trees that can be either cell lineage trees, binary trees where the cells form the leaves and mutations occur along tree branches, or mutation trees (Figure 1b) that depict the partial temporal order in which mutations were acquired [34]. For cell lineage trees a heuristic has been proposed for clustering cells into clones in a post-processing step [29] which results in trees that are closer to bulk clonal trees.

**Figure 1:**
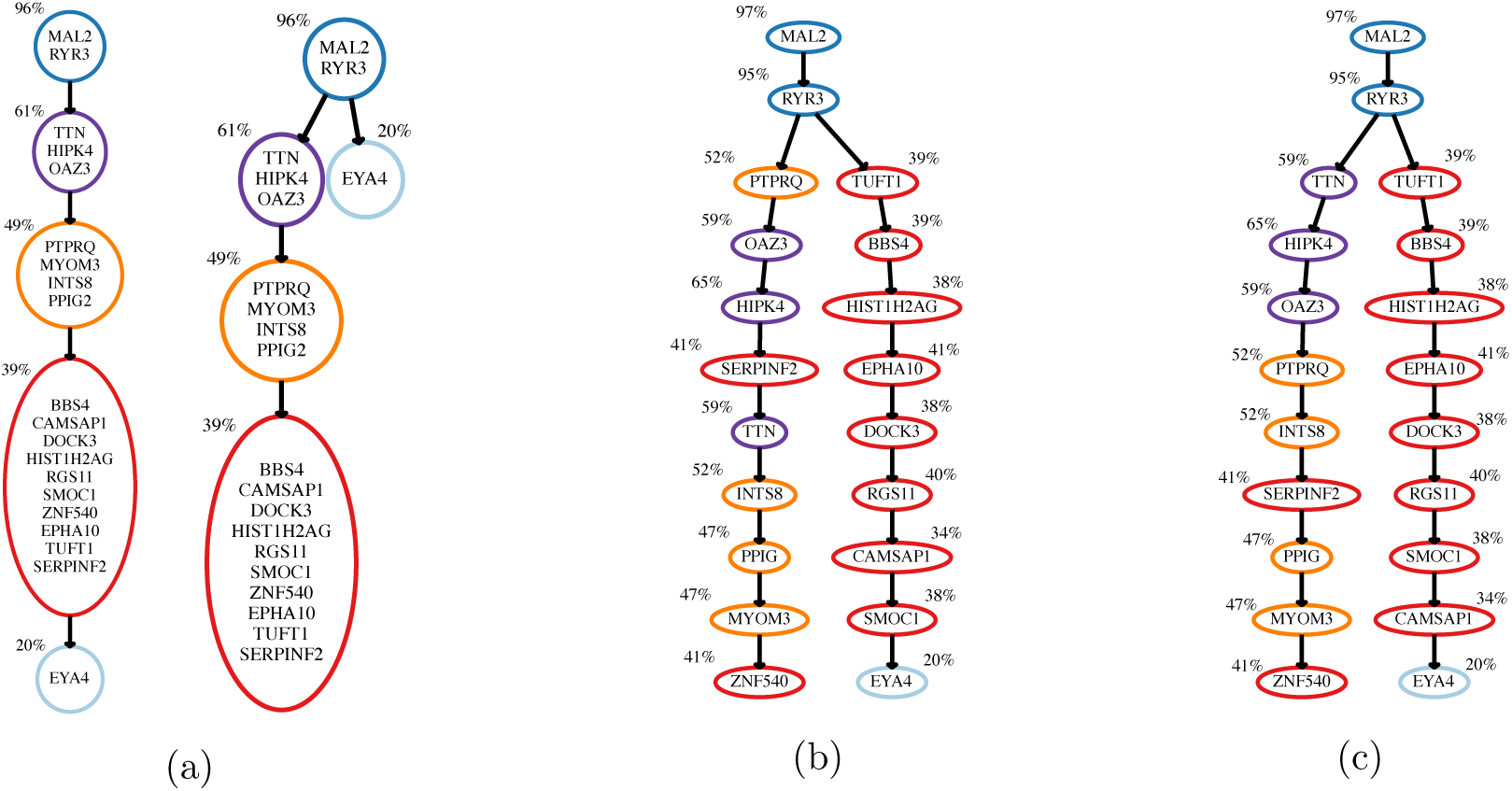
Mutation histories inferred from bulk and single-cell sequencing data for a patient with childhood ALL (Patient 1 of the study in [33]): (a) Clonal trees compatible with the bulk sequencing data (inferred with CTPsingle [20]). Clones are annotated with the mean VAF of the mutations newly acquired by the clone; (b) Mutation tree inferred with SCITE [28] from single-cell panel sequencing data of 111 cells. The colouring scheme follows from the clustering in the CTPsingle trees. Mutations are annotated with the cellular frequencies in the bulk sample. (c) Mutation tree inferred with B-SCITE from the combined single-cell and bulk data. B-SCITE infers the same early branching event as SCITE, but finds a mutation ordering that is in better congruence with the bulk frequencies.

Another difference is that the VAFs obtained from a bulk sample, are well suited for inferring the temporal order of mutations (by ordering mutations by decreasing VAF), but of limited use for the identification of branching events. On the contrary, single-cell genotypes obtained from SCS data have a lot of strength to infer branching events, but due to unobserved ancestral states and high noise levels may give ambiguous or no signals on the temporal order of mutations in linear segments of the inferred mutation trees.

As the strengths and weaknesses of single-cell and bulk data are to a large extent complimentary with respect to phylogeny inference, using both data types for a joint inference should improve our understanding of subclonal tumour evolution over using each data alone. It has already been shown that clustering of mutations into subclones from bulk sequencing data can be informed by single-cell genotypes to obtain more accurate results [35]. In this work, we present B-SCITE, a probabilistic approach for the inference of tumour phylogenies from combined single-cell and bulk sequencing data. We show that B-SCITE systematically outperforms competing approaches in terms of tree reconstruction accuracy on a comprehensive set of simulated datasets and provides more plausible mutation histories for real tumour data.

## 2 Methods

### 2.1 Tree models

A clonal tree *T* is a rooted tree with node set *V*(*T*) = {*υ*_1_,*υ*_2_, …, *υ_s_*,*υ_s_*_+1_} and root *υ_s_*_+1_ representing the population of healthy cells of a patient. The other nodes represent tumour subclones of the same individual which are all genetically different. This tree can be represented by an *s* × *s* ancestor matrix *A_T_* defined as follows:

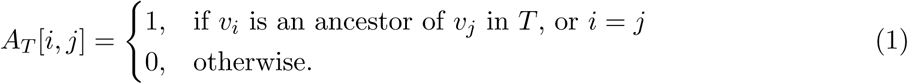

Let **M** = {*M*_1_, *M*_2_,…, *M_n_*} represent the set of somatic mutations detected in the tumour sample. We assume in the following that each mutation was acquired exactly once by one subclone and then passed on to all its descendants (infinite sites assumption). The function *N_T_*: **M** → *V*(*T*) \ {*υ_s_*_+1_} assigns the mutations to tree nodes, so that *N_T_*(*Mi*) denotes the subclone where mutation *M_i_* was acquired. As each subclone is genetically distinct, *N_T_* is required to be surjective onto *V*(*T*) \ {*υ_s_*_+1_}. Each non-root node *υ* can then be labelled with a mutation set *L_T_*(*υ*) = {*M_i_* ∣ *N_T_*(*M_i_*) = *υ*} = ∅. Then the set of mutations present in a subclone *υ_j_* ∊ *V*(*T*) \ *υ_s_*_+1_ consists of all mutations acquired along the path from the root to *υ_j_*, while the root node *υ_s_*_+1_ representing the population of healthy cells has no mutations. Each node *υ_i_* ∊ *V*(*T*) can be annotated with the relative frequency of its cell population in the analysed tumour.

A mutation tree can be considered as a special case of a clonal tree where *s* = *n* and each non-root node has a single and unique mutation as label, such that *N_T_* becomes a bijection. To simplify notation, we can always without loss of generality index the nodes of mutation tree such that *N_T_*(*M_i_*) = *v_i_*.

### 2.2 Input data

We assume a set M = {*M*_1_, *M*_2_, …, *M*_n_} of heterozygous somatic single-nucleotide variants from diploid regions of the genome is given. These mutations were observed in a tumour via sequencing of a bulk and multiple single-cell samples.

The bulk sequencing provided for each mutation *M_i_* the number of variant and total reads, denoted respectively by *r_i_* and *t_i_*, spanning the genomic position of *M_i_*. SCS provided the observed mutation profiles of m sequenced cells as the column vectors of a mutation matrix *D_n_*_×_*_m_* defined as

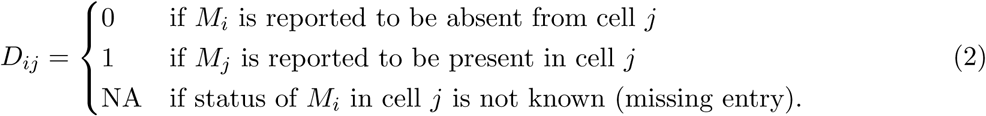

### 2.3 Tree scoring based on bulk sequencing data

Assume we are given a mutation tree *T* over *n* = *s* mutations whose (*s* + 1) nodes represent the set of cellular subpopulations in the analysed tumour. Our goal is to assign non-negative real values Φ = {*ϕ*_1_, *ϕ*_2_, …, *ϕ_s_*_+1_} to the nodes of *T* such that

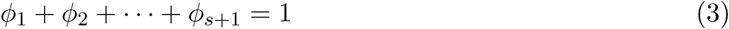

and the likelihood of bulk sequencing data (defined below) be maximised. Intuitively *ϕ_i_* represents the inferred fraction of the cellular population corresponding to node *υ_i_*. The inferred fraction *y_i_* of cells harbouring mutation M_i_, appearing for the first time at node *υ_i_* and being present at *υ_i_* and all of its descendants (i.e., nodes *υ_j_* such that *A_T_*[*i,j*] = 1), is then

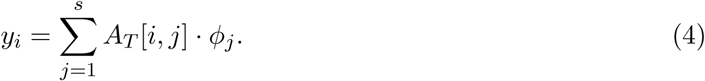

Alongside the tree constraints in (Equations 3 and (4, we introduce a likelihood model for the bulk sequencing data to allow us to combine it with the single-cell measurements.

Consider an arbitrary mutation *M* ∈ **M** with *t* total and *r* variant reads spanning its genomic position. If the underlying fraction *y* of cells from the sequenced bulk sample possesses this mutation, we assume that the sampled number of variant reads *r* follows a binomial distribution with parameters *t* (number of trials) and 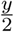 (success probability). Here, division by 2 is required since mutation *M* is heterozygous. For high coverage *t*, the binomial distribution can be approximated by the normal distribution with mean 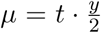 and standard deviation 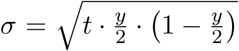. The logarithm of the probability density is

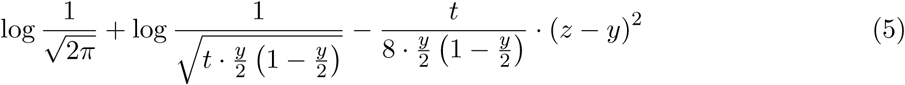

where 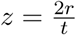, which represents the bulk sequencing data-based fraction of cells harbouring mutation *M*, based on the assumption that *M* is from a diploid region.

For a given assignment of values *y_i_*, after discarding constant terms, the log-likelihood of the entire bulk data is then

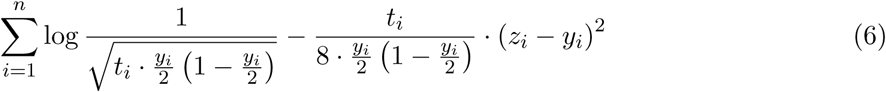

Our goal is to maximise the likelihood over the latent variables *yi,* under the constraints (3) and (4) imposed by the tree topology. To make this maximisation problem tractable by existing solvers, we use bulk data-derived frequencies 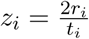 to approximate the standard deviations,

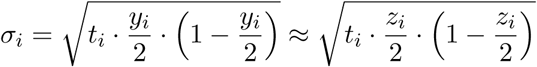

so that the log terms in Equation 6 become constant and can be removed from the optimisation. Our problem then transforms to maximising the likelihood over the underlying frequencies. We therefore define the score of the bulk data to be

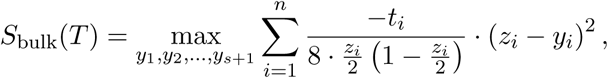

subject to the restrictions imposed in Equation 3 and Equation 4 where the sum involves the weighted quadratic terms from the Gaussian approximation. Obtaining the optimal values for *y_i_* represents an instance of a Quadratic Program (QP). After introducing additional variables in analogous ways as in [19], it reduces to a QP instance solvable by the existing commercial and free QP solvers. For our purposes, we have used IBM ILOG CPLEX Optimization Studio V12.5.

### 2.4 Tree scoring based on single-cell data

For the tree scoring based on single-cell data, we need to assess how well the observed mutation states of the single cells match the subclones defined by *T*. Due to noise in the mutation matrix *D*, the single cells will likely fit to none of the *s* + 1 cell populations defined by *T* perfectly, even if the tree represents the true mutation history of the tumour. We account for this by using the probabilistic error model introduced in [28]: Let the vector ***σ*** = (***σ***_1_, ***σ***_2_,…, ***σ****_m_*) define the attachments of the single cells to *T*, such that *s_j_*, the single cell corresponding to column *j* in *D*, attaches to *υ_σj_*. Then we expect *S_j_* to have the mutations assigned to the nodes Vj that belong to the path from root to *υ_σj_* (i.e. all nodes Vj such that *A_T_* [*i*, ***σ****_j_*] = 1). The observational probabilities are then

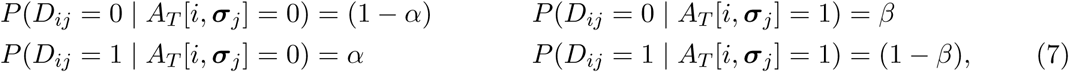

where *α* denotes the probability of observing a false positive and *β* denotes the probability of observing a false negative. The two error rates are summarised as *θ* = (*α*, *β*) in the following. By setting the probability of having missing observations (*D_ij_* = NA) to 1 independent of the true state, they are neutral and do not contribute to the tree scoring,

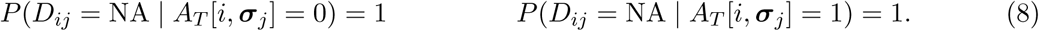

Assuming the observational errors are independent of each other, the likelihood of a given mutation tree *T*, sample attachment vector ***σ***, and *θ* is then

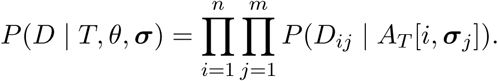

In the following we marginalise out the sample attachments to focus on the mutation tree as the informative part of the mutation history which is also more robust against noise than the location of individual samples in the tree. This give us

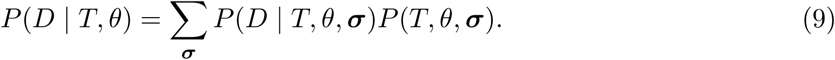

In practice SCS data is often contaminated with doublet samples. Therefore we modify our model such that each sample is treated as a weighted mixture of a singlet and doublet sample as previously described in [27]. The marginalised likelihood then becomes

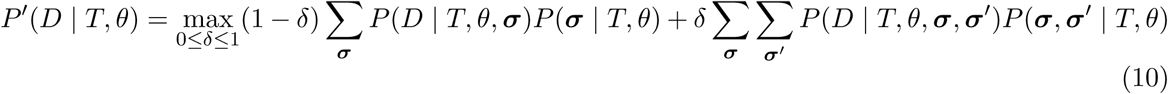

where *δ* is the probability of a sample being a doublet. The double sum over the attachment vectors ***σ*** and ***σ***′ creates all combinations of attachment pairs. This can be efficiently computed in time *O*(*mn*^2^), as the score can be computed separately for each sample as shown in [27].

Finally to obtain a single-cell based tree score that is on the same scale as the bulk score, we take the log of the marginalised likelihood,

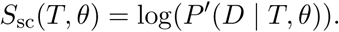

### 2.5 Combined B-SCITE approach

To measure how well a candidate mutation tree *T* fits a combination of bulk and single-cell measurements, we use the joint log-likelihood score defined as

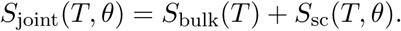

Our goal is to find a combination (*T*, *θ*)* that maximises the above score:

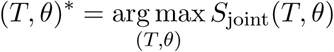

The number of possible mutation trees is too large to allow for an exhaustive search. Therefore we use a variant of the Markov chain Monte Carlo approach introduced in [28] to search the joint (*T, θ*) space. In each step it proposes a new state with proposal probability *q*(*T′*,*θ′* ∣ *T*, *θ*). The new state has either a new mutation tree *T′* or new parameters *θ′* and is accepted with probability

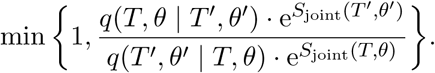

Note that *S_bulk_*(*T*) does not depend on *θ*. Therefore it needs not to be recomputed to obtain Sjoint (*T*, *θ′*) after a *θ*-move. A detailed description of the MCMC moves can be found in [28].

### 2.6 Compression of mutation trees into clonal trees

To compare the mutation trees inferred by B-SCITE to clone trees or mutational clusters, we cluster mutations along each linear chain of mutations between adjacent branching points in the mutation tree. We employ a 1D Gaussian mixture model with *K* components for the cellular frequencies *z_i_* of the mutations in the chain. The cluster assignments are learnt with the EM algorithm. The only difference from the standard approach is that the standard deviation of each mixture component is fixed by its mean *y_k_* according to (Equation 5 and the binomial approximation. The optimal number of mixture components is selected according to the AIC.

## 3 Results

We tested B-SCITE on a comprehensive set of simulated data. We focused initially on comparing with ddClone [35] which is, to the best of our knowledge, the only existing method that performs analysis of intra-tumour heterogeneity by jointly using bulk and SCS data. ddClone was shown to significantly outperform methods only utilising bulk sequencing data. For ease of comparison, we followed the simulation strategy employed in the original ddClone publication (see Section A.1 for details).

Since B-SCITE infers the phylogeny of the tumour’s mutations, we additionally compared to two tree inference methods, OncoNEM [29] and SCITE [28], working on single-cell data only.

## 3.1 Performance assessment on simulated data

### 3.1.1 V-measure comparisons of clustering accuracy

Since ddClone does not provide any output related to the tree of tumour evolution, we compared results using the V-measure of cluster assignments [36]. To process the results of B-SCITE into clusters, we clustered mutations along each linear chain in the inferred tree between each pair of consecutive branching points (see Section 2.6). OncoNEM also provides an option to cluster the data into subclones on the inferred phylogeny, and is hence included in the comparison.

The results (Figure 2 for 25 single cells and 10 clones) show that B-SCITE consistently outperforms ddClone. Both approaches are also robust to the doublet rate and the distortion in the single-cell data. OncoNEM, which only utilises the single-cell data, improves as the sampling more closely reflects the bulk tumour (as λ increases). For highly distorted data OncoNEM performs somewhat worse than ddClone, before performing notably better as the single-cell quality increases. Increasing the number of cells from 25 to 50 and 100 has a marginal effect on the accuracy (Figure A1, although more simulated clones may also be observed with more cells). However when simulating a smaller number of clones (Figure A2 with 6 clones instead of 10), OncoNEM outperforms ddClone even at the highest distortion (λ = 1) while B-SCITE remains the best performer.

**Figure 2:**
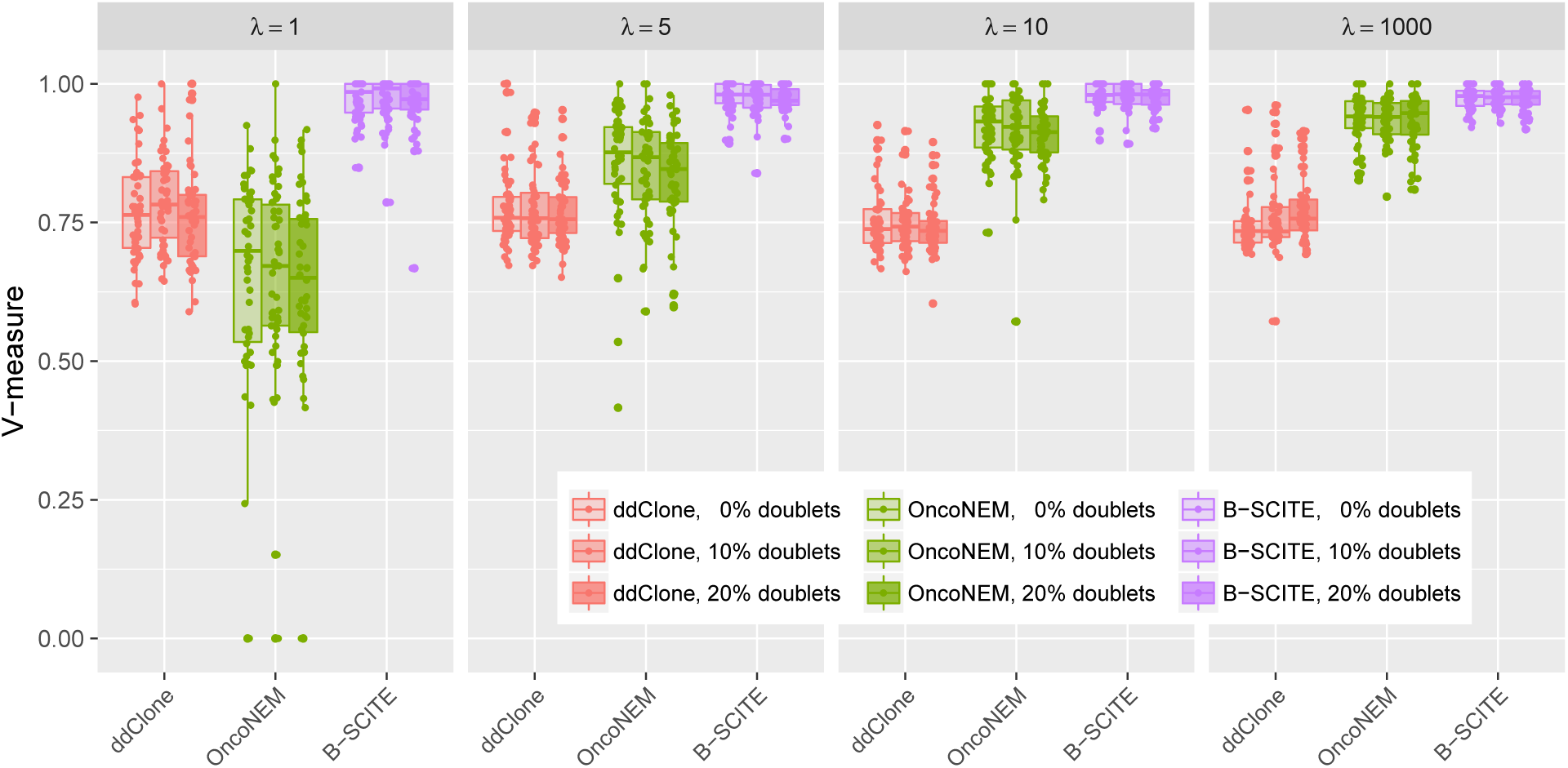
Comparison of inferred mutation clusters for ddClone, OncoNEM and B-SCITE. The simulated data consists of 10 clones, from which 25 cells are sampled with a false negative rate of 0 .2. Simulations are run for various doublet rates and values of the distortion parameter λ which modifies the single-cell frequencies away from the bulk frequencies.

**Figure A1:**
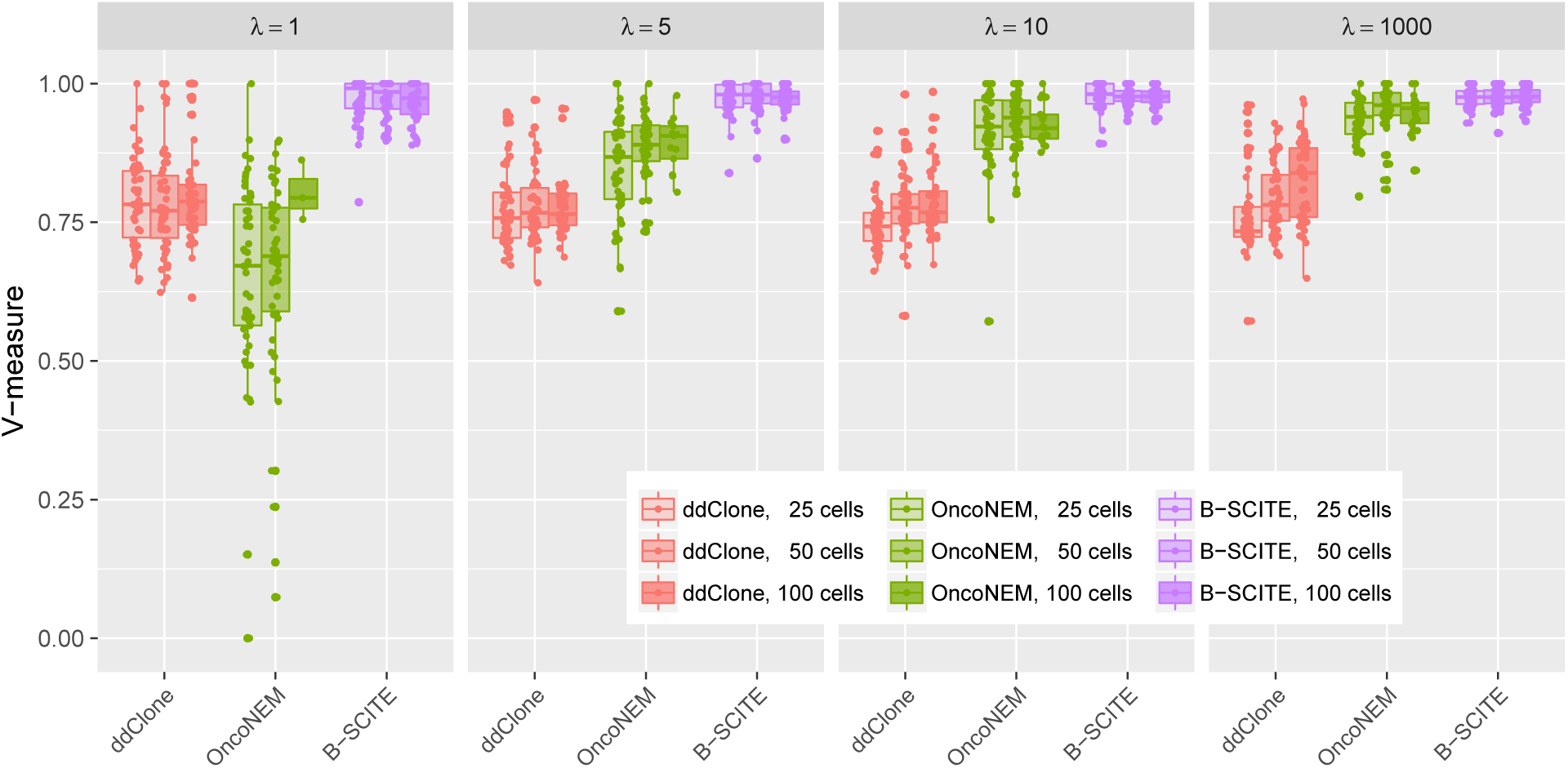
Comparison of inferred mutation clusters when 25, 50 or 100 single cells are sampled for ddClone, OncoNEM and B-SCITE. The simulated data consists of 10 clones with a doublet rate of 0.1 and a false negative rate of 0.2.

**Figure A2:**
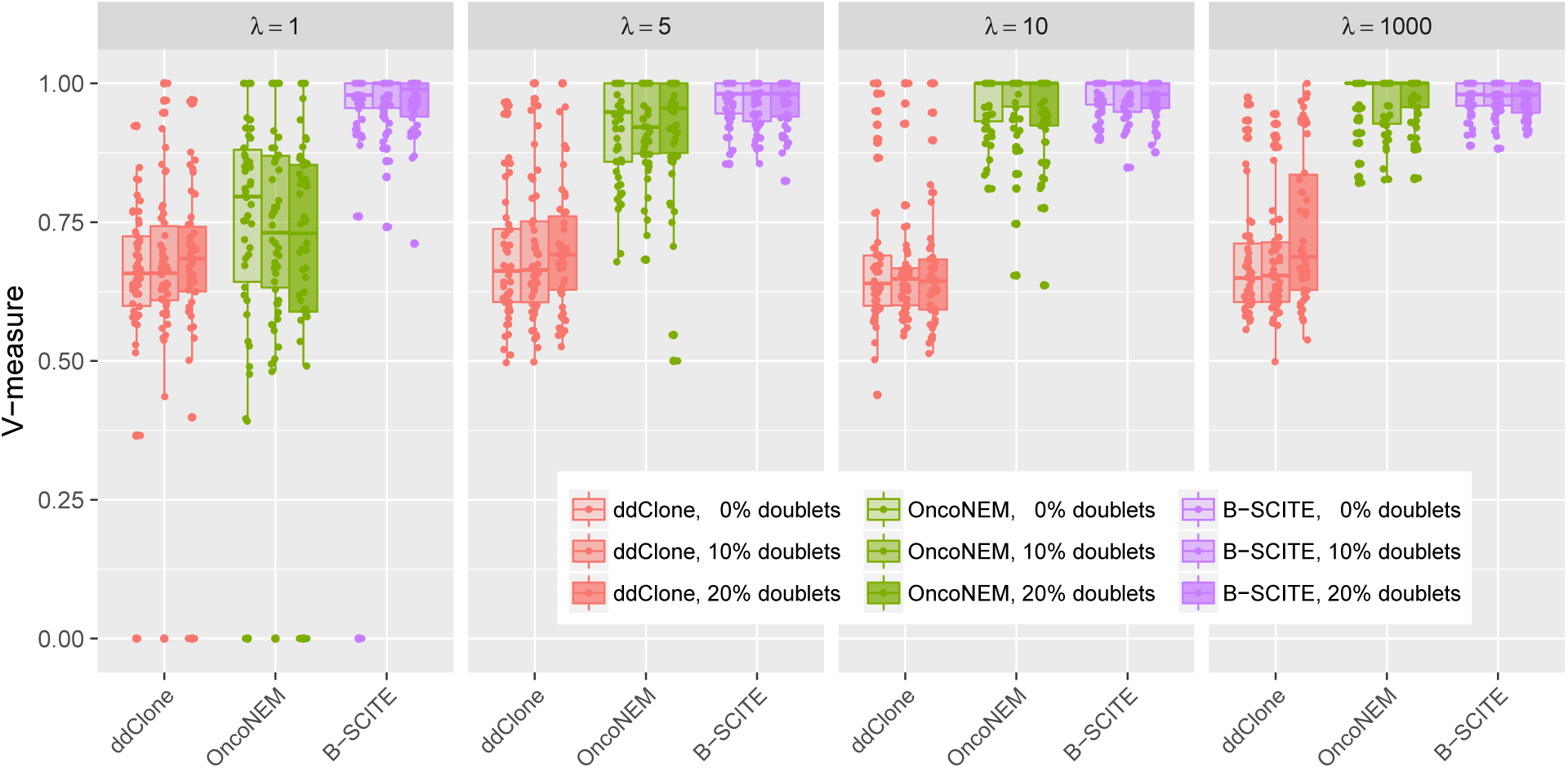
Comparison of inferred mutation clusters when 6 clones are simulated for ddClone, OncoNEM and B-SCITE. 25 cells are sampled from the 6 clones with a false negative rate of 0.2.

### 3.1.2 Accuracy in inferring phylogenetic order of mutations

As well as outperforming ddClone on clustering a tumour’s mutations (Figure 2), B-SCITE also infers the complete phylogenetic history of the tumour. We therefore compared B-SCITE to the single-cell phylogenetic methods OncoNEM and SCITE. Specifically, for SCITE we chose the extended version with the doublet model [27] to make sure that any change in performance can be fully attributed to the additional data available to B-SCITE. We considered three distance measures between the inferred and ground truth trees (Section A.2).

B-SCITE again has the best and most robust performance over the range of λ (Figure 3) with both single-cell methods improving as the single-cell sampling approaches the bulk frequencies. The apparent improvement for B-SCITE as λ decreases is due to smaller number of observed clones being included in the accuracy measure.

**Figure 3:**
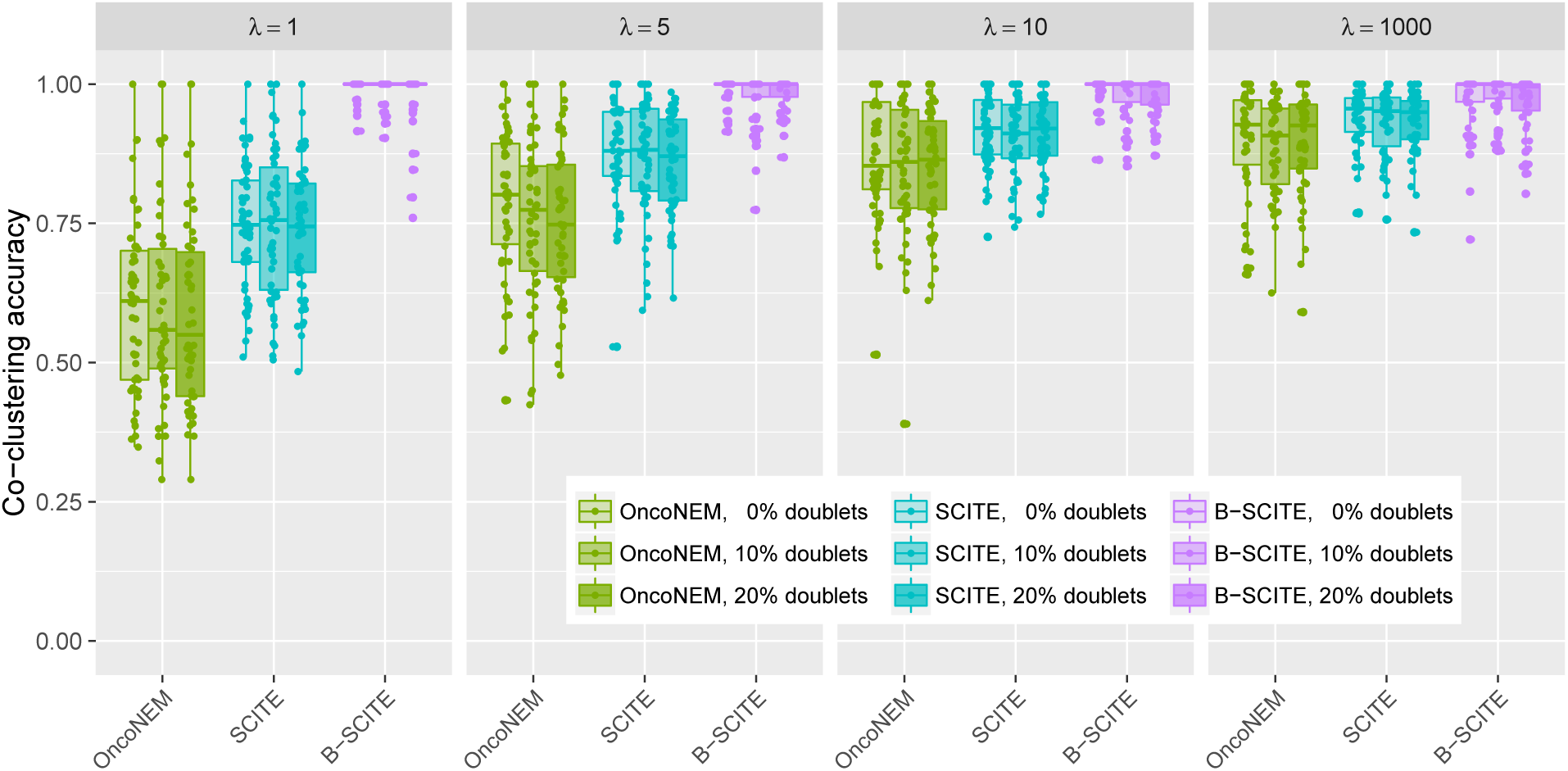
Comparison of phylogenetic inference for OncoNEM, SCITE and B-SCITE. The simulated data is identical to Figure 2 and consists of 10 clones, from which 25 cells are sampled with a false negative rate of 0.2.

Changing the number of cells (Figure A3) or the number of clones (Figure A4) we observe the same pattern of SCITE performing slightly better than OncoNEM, and with B-SCITE on top. Similar behaviour is also observed for inferring the correct ancestry relationships (Figure A5). For mutations in separate lineages, both SCITE and B-SCITE perform well in correctly detecting the correct separations, while OncoNEM has somewhat worse accuracy (Figure A6).

**Figure A3:**
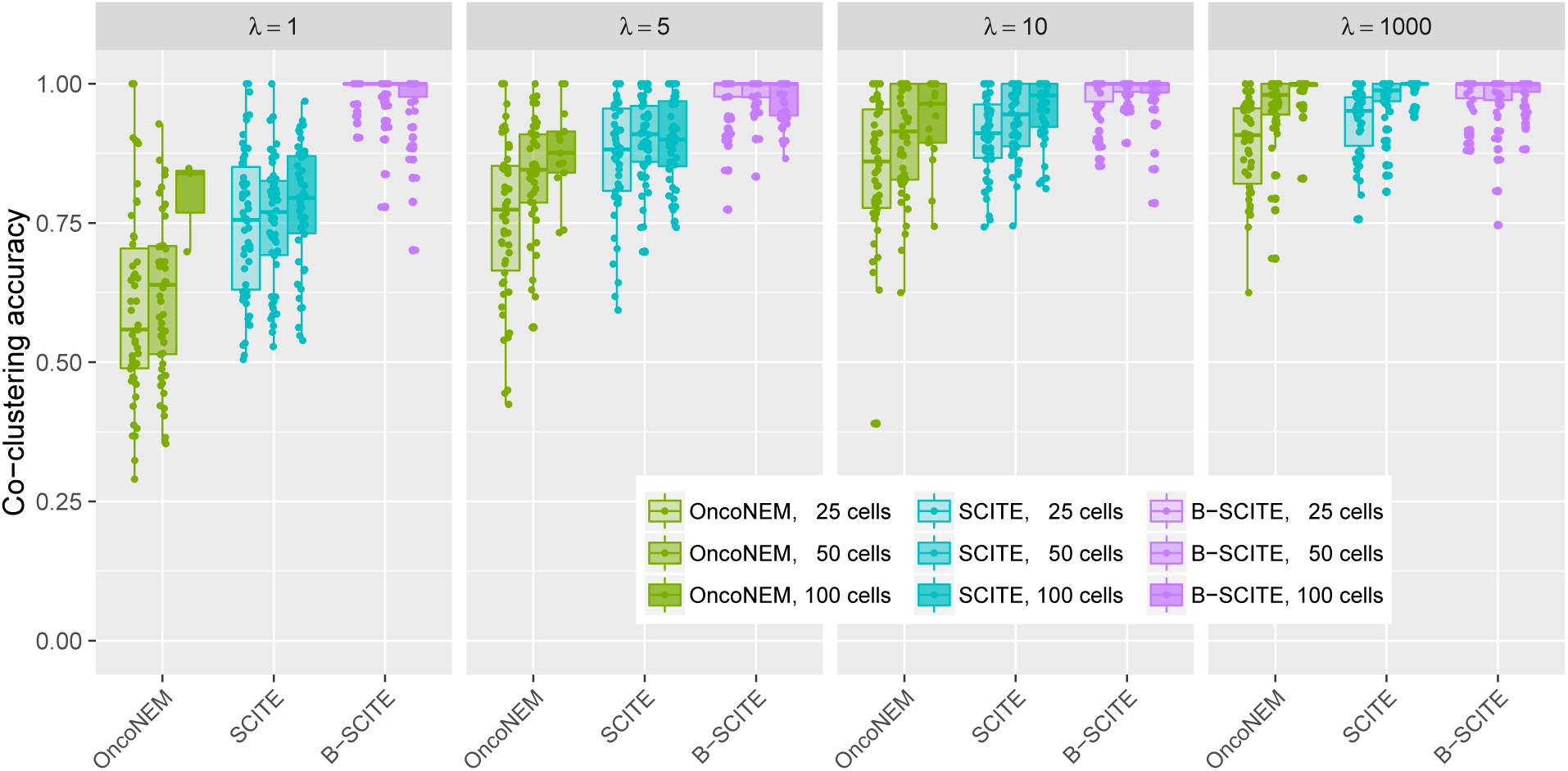
Comparison of phylogenetic inference when 25, 50 or 100 single cells are sampled for OncoNEM, SCITE and B-SCITE. The simulated data is identical to Figure A1 with 10 clones, a doublet rate of 0.1 and a false negative rate of 0.2.

**Figure A4:**
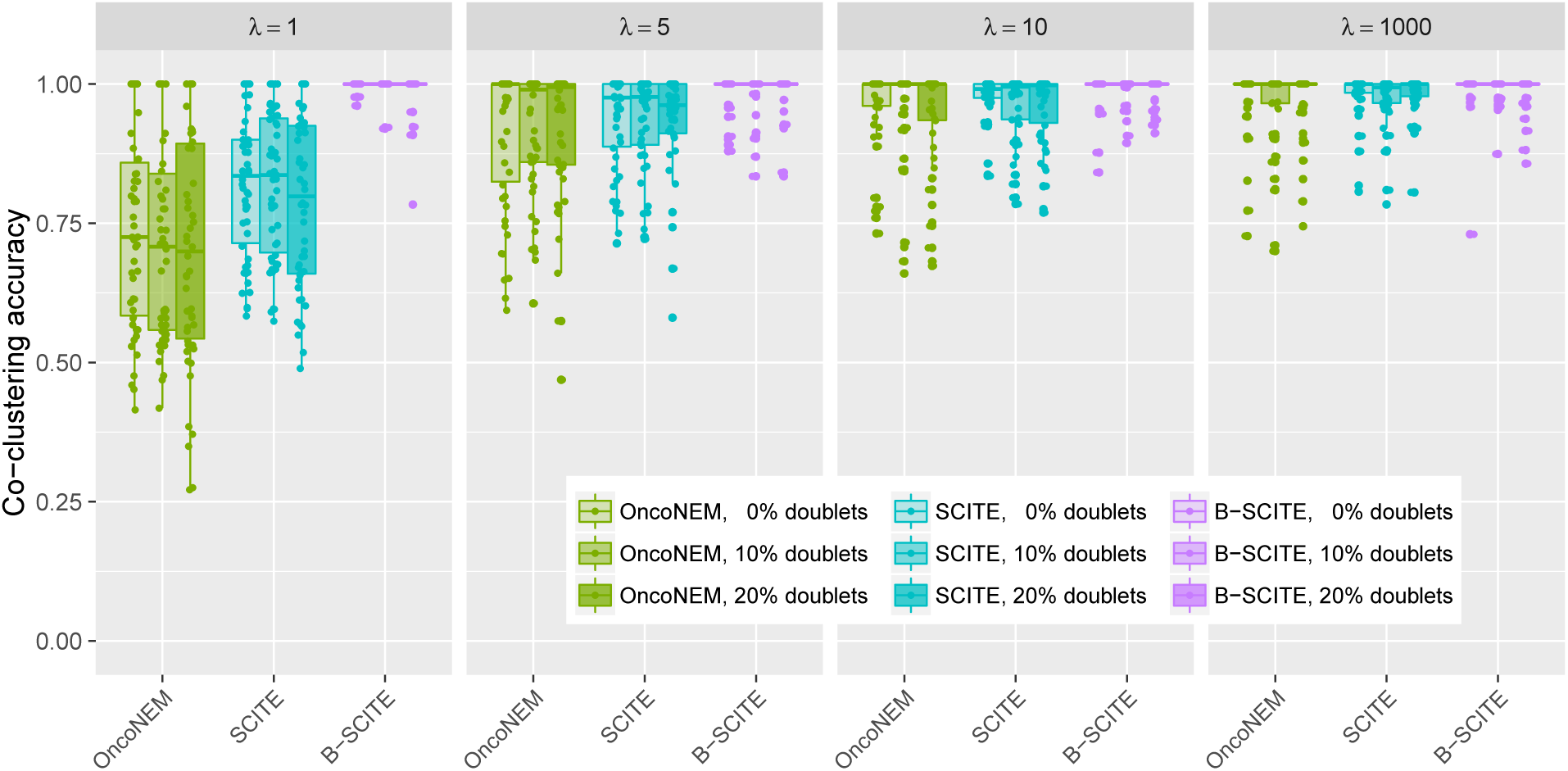
Comparison of phylogenetic inference when 6 clones are simulated for OncoNEM, SCITE and B-SCITE. The simulated data is identical to Figure A2 with 25 cells and a false negative rate of 0.2.

**Figure A5:**
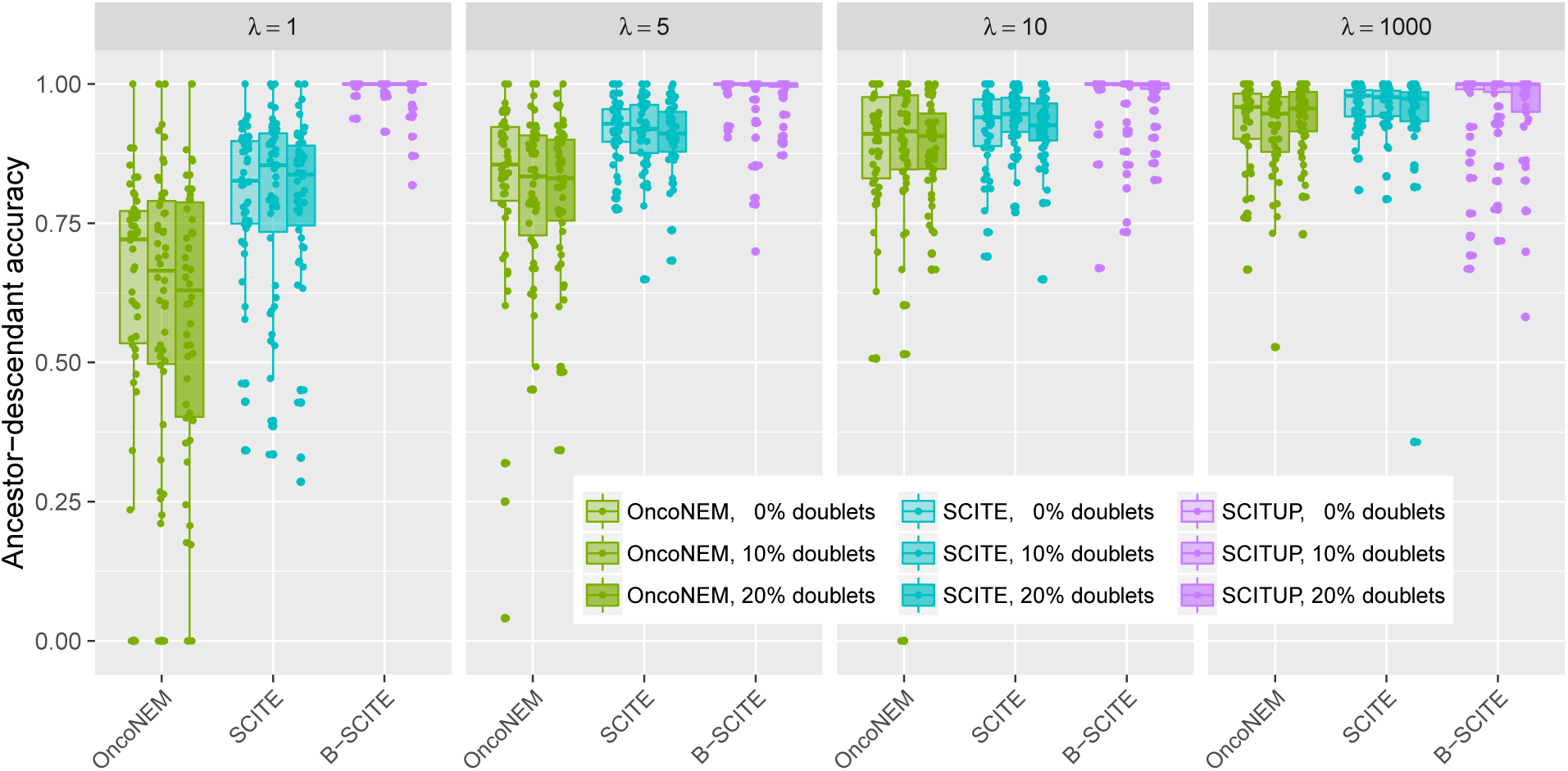
Comparison of ancestry relationship inference for OncoNEM, SCITE and B-SCITE. The simulated data is identical to Figure 2 with 10 clones, 25 cells and a false negative rate of 0.2.

**Figure A6:**
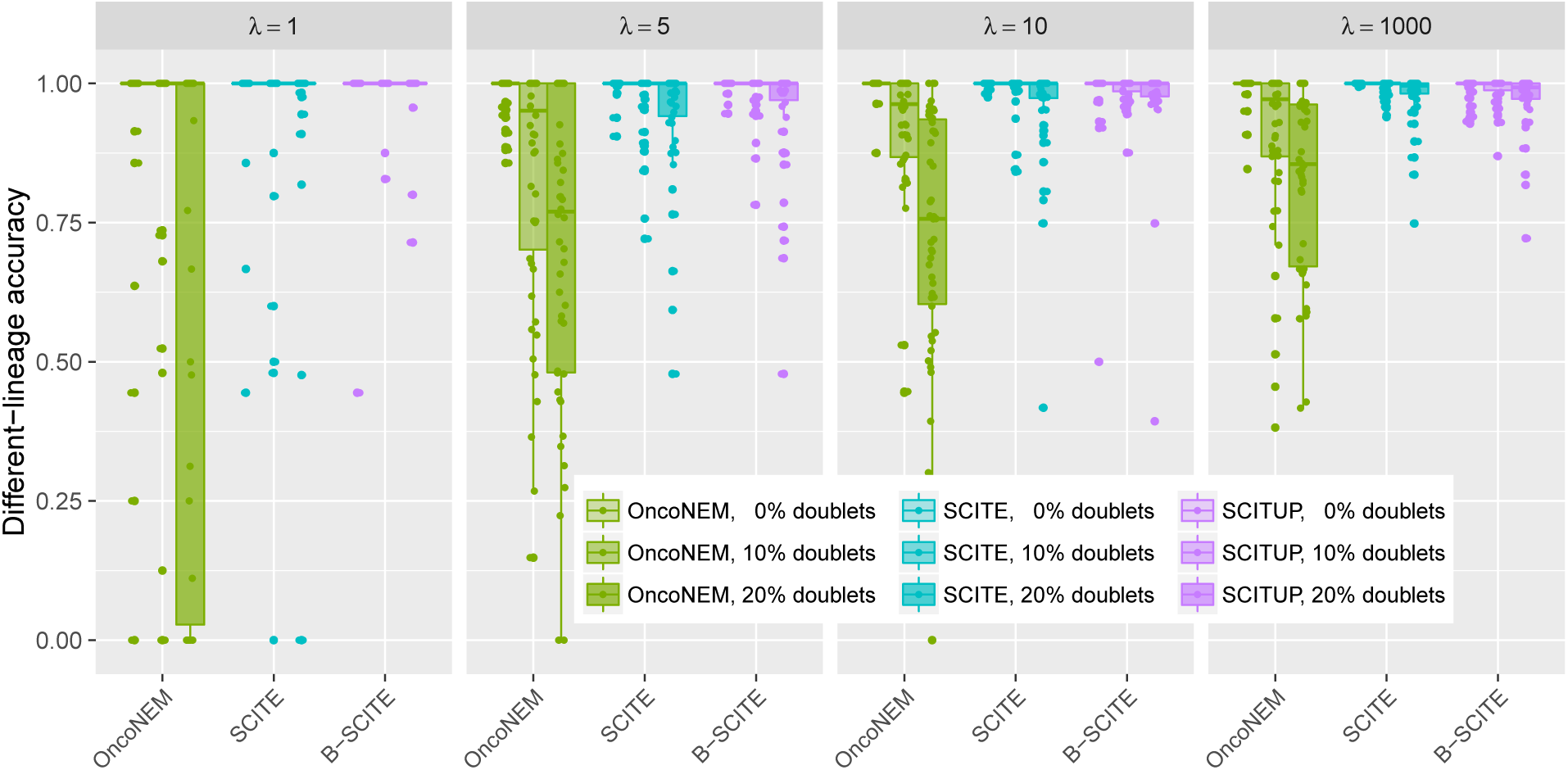
Comparison of lineage separation inference for OncoNEM, SCITE and B-SCITE. The simulated data is identical to Figure 2 with 10 clones, 25 cells and a false negative rate of 0.2.

### 3.2 Application to real data

To assess the performance of B-SCITE on real tumour data, we analysed the sequencing data of two patients with childhood leukaemia previously published in [33]. For both patients, a bulk sample was sequenced together with a large number (> 100) of single cells for which targeted sequencing was performed using a personalised panel. This allows us to compare B-SCITE with methods relying solely on single-cell or bulk sequencing data. For our comparison, we chose CTPsingle [20] for the bulk-only approach and SCITE [28] for single-cell based inference.

For patient 1, 20 mutations were detected sequencing one bulk sample and 111 single cells. The phylogenies inferred by B-SCITE and its two competitors are depicted in Figure 1. CTPsingle, the approach based on bulk samples, finds two trees compatible with the observed frequencies. Both trees cluster the 20 mutations in five subclones and are either completely linear or have a single mutation in a separate branch. Using the SCS data of the same patient SCITE detects an early branching event that splits up some of the subclones inferred by CTPsingle. Without knowledge of the ground truth tree there is no certainty whether the branching reflects the true phylogeny. However, having data from such a large number of cells and the close location to the root, makes it highly likely that the branching is genuine. The same branching is also inferred by B-SCITE which finds generally the same topology as SCITE, but arranges mutations differently in linear segments of the tree. As expected, having the additional information of VAFs for the individual mutations allows B-SCITE to find an ordering that is in better congruence with the observed mutation frequencies, which should be in decreasing order. As a consequence, the mutation ordering inferred by B-SCITE is also closer to the subclone clustering of CTPsingle, in the sense that mutations from the same cluster that are in the same branch tend to be closer together than in the SCITE tree.

For patient 2, 16 mutations were detected sequencing one bulk sample and 115 single cells. CTPsingle reports a subclone clustering that is compatible with many phylogenies and some of them are depicted in Figure A7. SCITE and B-SCITE each infer a single tree Figure 4(b)-(c). The general topologies of the two trees are again similar with different arrangement of mutations on linear segments. In particular, B-SCITE puts the mutations in *FAM105A* and *CMTM8* higher up in their branch which better reflects their relatively high frequencies (44% and 42%). Notably, B-SCITE does not completely sort the mutations in this branch by frequency as the mutation in *RRP8* is placed between two mutations with higher frequencies. This indicates a strong signal for this placement is coming from the single-cell data. Apart from issues with mutation calling, a possible explanation could be a copy number change affecting *RRP8* which decreases its observed VAF.

**Figure 4:**
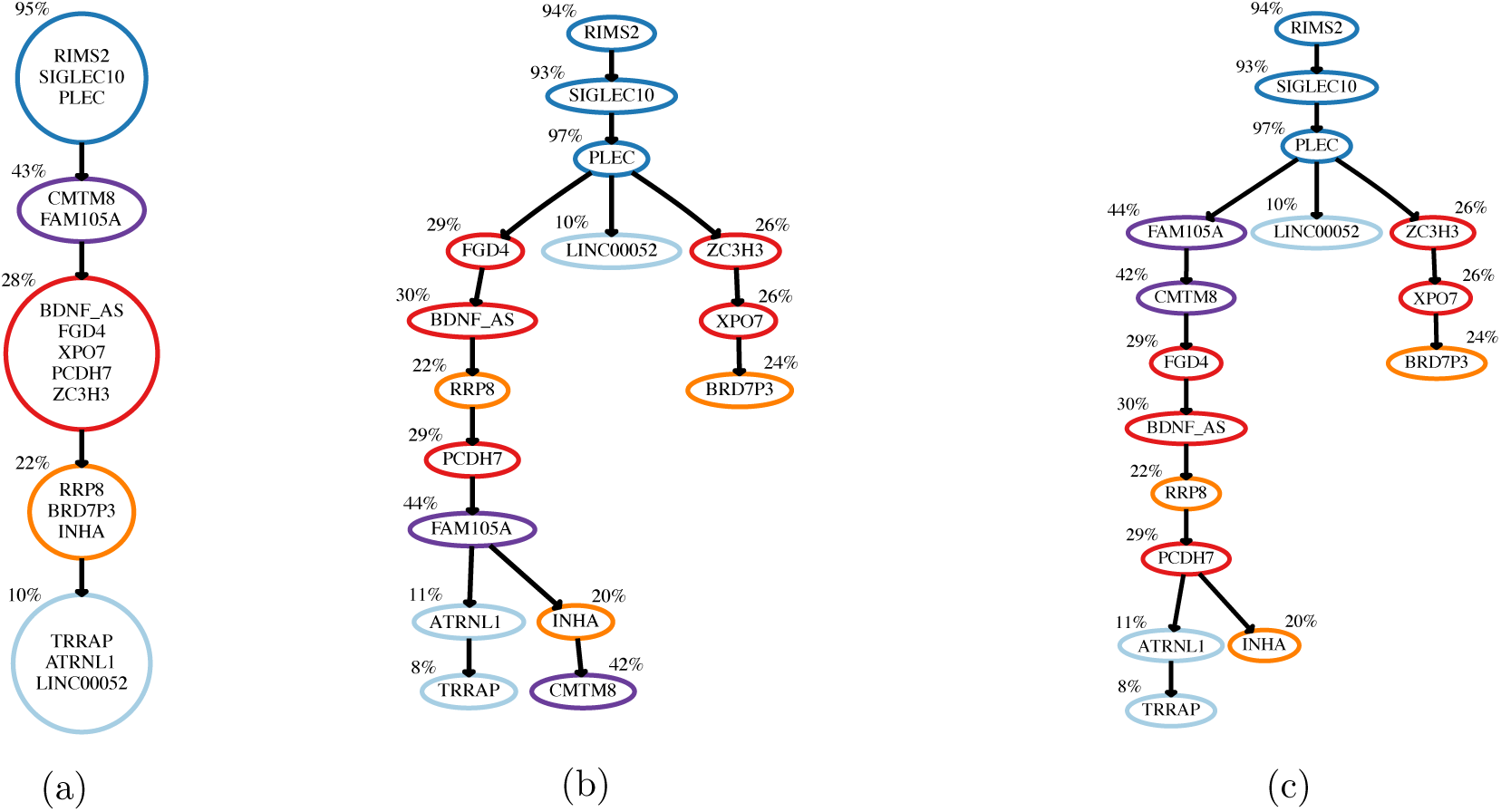
Mutation histories inferred from bulk and single-cell sequencing data for a patient with childhood ALL (Patient 2 of the study in [33]): (a) One of the multiple clonal trees compatible with the bulk sequencing data (inferred with CTPsingle [20]). Clones are annotated with the mean VAF of the mutations newly acquired by the clone; Several other compatible trees are depicted in Figure A7 (b) Mutation tree inferred with SCITE [28] from single-cell panel sequencing data of 115 cells. The colouring scheme follows from the clustering in the CTPsingle tree. Mutations are annotated with the cellular frequencies in the bulk sample. (c) Mutation tree inferred with B-SCITE from the combined single-cell and bulk data.

**Figure A7:**
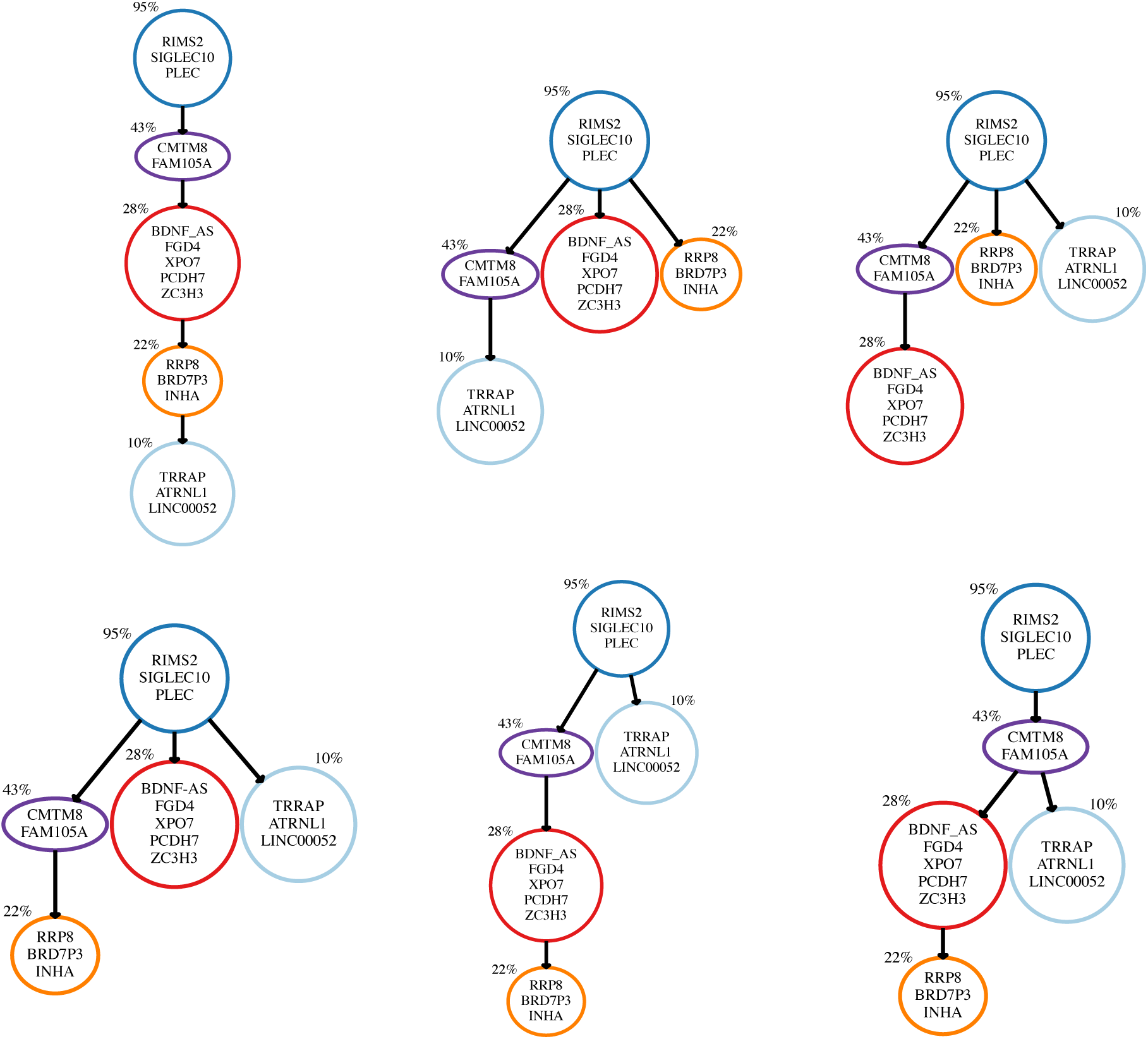
Subset of clonal trees inferred with CTPsingle for patient 2 of the leukaemia dataset in [33]: The clonal frequencies of the five clusters are compatible with multiple different tree topologies and for some topologies multiple optimal assignments of clusters to nodes exist.

## 4 Conclusions

Recent advances in sequencing technologies allow large scale bulk and single-cell sequencing of tumour samples. The resulting data is invaluable for understanding of the evolutionary history and subclonal composition of individual tumours and addressing the issue of treatment failure due to resistant cell populations. The bottleneck to fully leveraging the joint strength of both data types is the lack of specialised software which integrates single-cell and bulk sequencing data in a joint inference scheme. Prior to this work, only a single tool (ddClone) has been available where subclone inference based on bulk sequencing data is informed by single-cell genotypes, but no integrative tool has been published for phylogeny inference. To fill this gap, we have developed B-SCITE, the first approach for inferring tumour phylogenies and subclonal compositions from combined single-cell and bulk sequencing data. B-SCITE uses a joint likelihood model to integrate both datatypes and performs a probabilistic search to find the best combination of a fully-resolved mutation history and values for the model parameters. Extensive simulation studies show that B-SCITE systematically outperforms competing single-cell based approaches thereby indicating that bulk data makes a valuable contribution to the inference. The quantity of the performance gain depends on the degree of sampling distortion between the single-cell samples and the composition of the bulk tumour. However, even in cases where the single-cell data very well reflect the tumour composition, B-SCITE outperforms its competitors.

To compare B-SCITE with ddClone, the only other tool combining single-cell and bulk sequencing data, we obtained subclones from B-SCITE’s fully-resolved mutation histories by performing a local mutation clustering. Experiments on a comprehensive set of simulated datasets showed that B-SCITE systematically outperforms ddClone suggesting that subclone inference benefits from taking the underlying phylogeny into account.

In addition, we explored the usefulness of combining bulk and single-cell data in the analysis of real tumours. Looking at the data from two patients with childhood leukaemia, we find that B-SCITE and the single-cell only approach SCITE infer very similar branchings in the tumour phylogenies, while B-SCITE shows an improved ability in reconciling the temporal ordering of mutations with their VAFs. In cases were B-SCITE refrains from ordering mutations by decreasing VAFs, we suspect the presence of a copy number change whose signal was overridden by a strong contrary signal in the single-cell data. In this sense, B-SCITE appears to be to some extent robust against deviations from the limiting assumption of all mutations being heterozygous and at copy number neutral sites. However, to further improve our model it should be generalised to mutation patterns that do not follow the infinite sites assumption.

## A Appendix

### A.1 Generation of simulated datasets

Using the notation from Section 2.1 we randomly generated 50 clonal trees of tumour evolution for 10 and 6 cell populations including a normal cell population (*s* = 9 and *s* = 5). For each of these 100 trees, we randomly spread 50 mutations to its nodes. In order to account for the population of healthy cells, we did not assign any mutation to the root node. Furthermore, to ensure that each node corresponds to a different subclonal population, unique with respect to the set of mutations it harbours, we assigned at least one mutation to each of the *s* non-root nodes.

While simulating frequencies *ϕ_i_* of cellular populations we set the lower bound for *ϕ_i_* to 0.02. To simulate values of *ϕ_i_* satisfying this constraint, we first simulated *s* + 1 random numbers *w*_1_, *w*_2_, …, *w_s_*_+1_ from the interval (0,1) and for each i = 1, 2, …, *s* + 1 assigned value *ϕ_i_* as

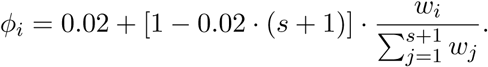

For mutation *M* with cellular prevalence *y* (that is calculated as the sum of frequencies of cellular populations harbouring *M*) bulk sequencing read counts were drawn from binomial distribution with parameters 10000 (number of trials) that accounts for sequencing coverage and was also previously used in [35] and 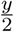 (success probability).

Before sampling single-cell data from the subclone genotypes, we first added sampling distortion to the subclone frequencies Φ to account for the fact that the single cell samples may come from a tumour region with a different subclonal composition than the bulk sample. The observed prevalences for each genotype are sampled from a Dirichlet distribution Φ*_observed_* ~ *Dir* (λΦ) as in [35]. The larger the value of λ the smaller is the difference between the observed frequencies and the bulk subclone frequencies.

To obtain the genotypes of the *m* single-cell samples, we then sampled *m* times (with replacement) from the *s* tumour subclone genotypes using the observed frequencies as sampling weights. We then converted each sample with probability *δ* into a doublet by sampling a second genotype and replacing the sample’s original genotype with the union of the two genotypes. The obtained genotypes form a *m* × *n* binary matrix. We then added false positives and false negatives by flipping each 0 in the mutation matrix to a 1 with probability *α* and each 1 to a 0 with probability *β*\* = *e^z^* · *β*, where *z* is sampled individually for each tree from a normal distribution with standard deviation 0.1 to obtain around 10% misspecification for the false negative rate. Finally, we added missing data points (NA) by removing each matrix entry with probability *μ*.

Only mutations present in the single cells we retained for analysis. For high distortion rates (low λ) of the single-cell sampling, some clones may not be present in any single cells. These clones are correspondingly removed from the ground truth tree when comparing the accuracy of different methods.

### A.2 Phylogenetic accuracy measures

Assuming that *T* and *I* represent simulated ground truth and inferred trees, respectively, the three measures used for tree comparison are defined as follows:

1. ***Ancestor-descendant accuracy:*** For this accuracy measurement, we consider all pairs of mutations (*a, b)* that are in ancestor-descendant relation in *T*. In other words, *A*[*N_T_*(*a*),*N_T_*(*b*)] = 1 and *N_T_*(*a*) ≠ *N_T_*(*b*). For each pair, we check whether this dependency between *a* and *b* is preserved in *I*. The total score is defined as number of preserved relations in *I* among such mutation pairs divided by their total number in *T*.
2. ***Different-lineage (siblings) accuracy***: Here we consider only pairs of mutations (*a, b*) that belong to different lineages in *T*. In other words *A*[*N_T_*(*a*), *N_T_*(*b*)] = *A*[*N_T_*(*b*), *N_T_*(*a*)] = 0. Analogously to the previous measure, we count the number of times this relation is preserved in *I* and divide by the total number of pairs of mutations satisfying this relation in *T*.
3. ***Co-clustering accuracy***: It is expected that mutations originating from the same subclone in *T* lie on the same lineage and in close proximity in *I*. In this measure, for each pair of mutations (*a, b*) such that *N_T_*(*a*) = *N_T_*(*b*) we define the co-clustering score of pair (*a, b*) as 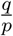, where *p* represents the total number of mutations on the path from *N_I_*(*a*) to *N_I_*(*b*), including the end-nodes, and *q* represents number of mutations *c* on this path such that *N_T_*(*c*) = *N_T_*(*a*). The total score of *I* is defined as the average of the scores over all mutation pairs belonging to the same subclone in *T*.

### A.3 Details of running ddClone, OncoNEM and SCITE

Below we provide brief description of input data and the main parameter settings used for running three methods that B-SCITE was compared against.

### A.3.1 ddClone

Genotype matrix, that is recommended input to ddClone, was obtained from single-cell data matrix *D* by the use of Single Cell Genotyper [37]. ddClone also requires purity value as part of the input. In each case, the exact simulated value of purity was provided. In each case, we run ddClone for 300 iterations. The choice of this value was primarily motivated by the default value set to 100 in the examples published with this tool that are of comparable size to our simulated datasets. In addition, running time of ddClone increases significantly as number of iterations is further increased.

### A.3.2 OncoNEM

The input to OncoNEM was directly obtained from matrix *D* after removing cells (i.e. columns) having none of the entries equal to 1, as such cells are assumed to be filtered prior to running this tool. For each run, values of false positive and false negative error rates were given as values of *α* and *β* used while simulating matrix *D* (see A.1).

### A.3.3 SCITE

Single-cell data input to SCITE is identical to matrix *D*, hence no input pre-processing was required. We run this tool for 3 repeats with 200 000 iterations for each repeat. Values of false positive and false negative error rates were provided analogously as for OncoNEM.

